# Metabolic basis of brain-like electrical signalling in bacterial communities

**DOI:** 10.1101/553305

**Authors:** Rosa Martinez-Corral, Jintao Liu, Arthur Prindle, Gürol M. Süel, Jordi Garcia-Ojalvo

**Affiliations:** Department of Experimental and Health Sciences, Universitat Pompeu Fabra, Barcelona Biomedical Research Park, 08003 Barcelona, Spain; Center for Infectious Diseases Research and Tsinghua-Peking Center for Life Sciences, School of Medicine, Tsinghua University, 100084 Beijing, China; Department of Biochemistry and Molecular Genetics, Feinberg School of Medicine, Northwestern University, Chicago, IL 60611, USA; Center for Synthetic Biology, Northwestern University, Evanston, IL 60208, USA; Division of Biological Sciences, San Diego Center for Systems Biology, and Center for Microbiome Innovation, University of California San Diego, California 92093, USA

**Keywords:** membrane potential, electrical signalling, potassium waves, bacterial biofilms, cellular excitability

## Abstract

Information processing in the mammalian brain relies on a careful regulation of the membrane potential dynamics of its constituent neurons, which propagates across the neuronal tissue via electrical signalling. We recently reported the existence of electrical signalling in a much simpler organism, the bacterium *Bacillus subtilis*. In dense bacterial communities known as biofilms, nutrient-deprived *B. subtilis* cells in the interior of the colony use electrical communication to transmit stress signals to the periphery, which interfere with the growth of peripheral cells and reduce nutrient consumption, thereby relieving stress from the interior. Here we explicitly address the interplay between metabolism and electrophysiology in bacterial biofilms, by introducing a spatially-extended mathematical model that combines the metabolic and electrical components of the phenomenon in a discretised reaction-diffusion scheme. The model is experimentally validated by environmental and genetic perturbations, and confirms that metabolic stress is transmitted through the bacterial population via a potassium wave. Interestingly, this behaviour is reminiscent of cortical spreading depression in the brain, characterised by a wave of electrical activity mediated by potassium diffusion that has been linked to various neurological disorders, calling for future studies on the evolutionary link between the two phenomena.

## 1 Introduction

Local interactions between individual cells are known to generate complex emergent behaviour in multicellular organisms, which underlie the functionalities of tissues, organs, and physiological systems [1]. The human brain provides one of the finest examples of such phenomena, with its complexity ultimately emerging from the interactions between neuronal cells [2]. Similarly, unicellular organisms can also self-organise into communities with complex community-level phenotypes [3]. Bacterial biofilms, in particular, provide a good model system for the study of collective behaviour in biological systems [4].

We have previously shown that biofilms of the bacterium *B. subtilis* can display community-level oscillatory dynamics [5]. These oscillations ensure the viability, under low-nitrogen (low-glutamate) conditions, of the cells in the centre of the community, which are crucial for community regrowth upon external sources of stress like antibiotics. In our experimental setup (Fig. 1A), growing two-dimensional biofilms that reach a certain size develop oscillations [6], such that peripheral cells periodically stop their growth (see black dotted line in Fig. 1B,C). This behaviour was attributed to a metabolic codependence between central and peripheral cells mediated by ammonium [5] (see introduction to Section 2 below).

**Figure 1:**
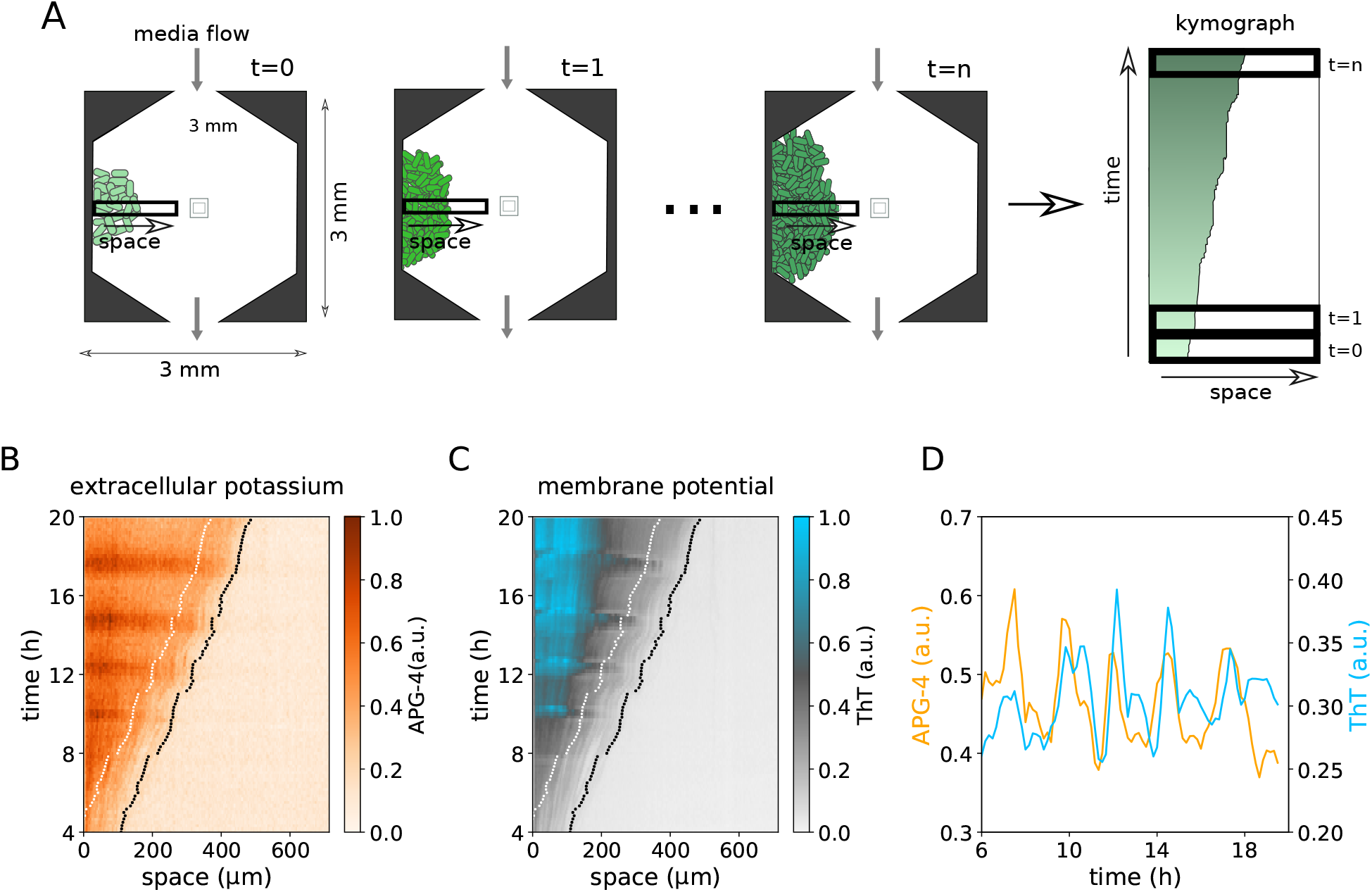
Biofilms of *B. subtilis* display oscillations in growth rate and electrical signalling activity. A) Scheme of the microfluidics device used to grow biofilms. Biofilms grow around the central pillar or attached to the wall, as in the cartoon. Biofilm dynamics can be represented as a kymograph, where each row represents a one dimensional cross-section of the biofilm area at a given movie frame. B-D) Experimental kymographs of a cross-section of a biofilm, as depicted in A, for different channels. Each pixel is the average of a region of 5.2 × 5.2 *μm*. The dotted black line denotes the limit of the biofilm as established from the phase-contrast images. B) Oscillations in extracellular potassium, reported by APG-4. C) Oscillations in membrane potential, reported by ThT. D) Extracellular potassium and ThT time traces corresponding to the position denoted by the white line in panels B,C. Data has been smoothed with an overlapping sliding window of size 3 frames.

Besides exhibiting metabolic oscillations in growth rate, our biofilms display periodic changes in extracellular potassium (Fig. 1B) and cellular membrane potential (Fig. 1C), as reported by the fluorescent dyes APG-4 and Thioflavin-T (ThT), respectively. The membrane potential changes reported by ThT are caused by the periodic release of intracellular potassium ions (Ref.[7] and Fig. 1D). Potassium propagates from the centre of the biofilm to its periphery according to the following scenario: potassium released by glutamate-deprived (and thus metabolically stressed) cells in the centre diffuses towards neighbouring cells, causing depolarisation, stress and subsequent potassium release (Fig. 2-right), which thereby keeps propagating along the biofilm in a bucket-brigade manner [7].

**Figure 2:**
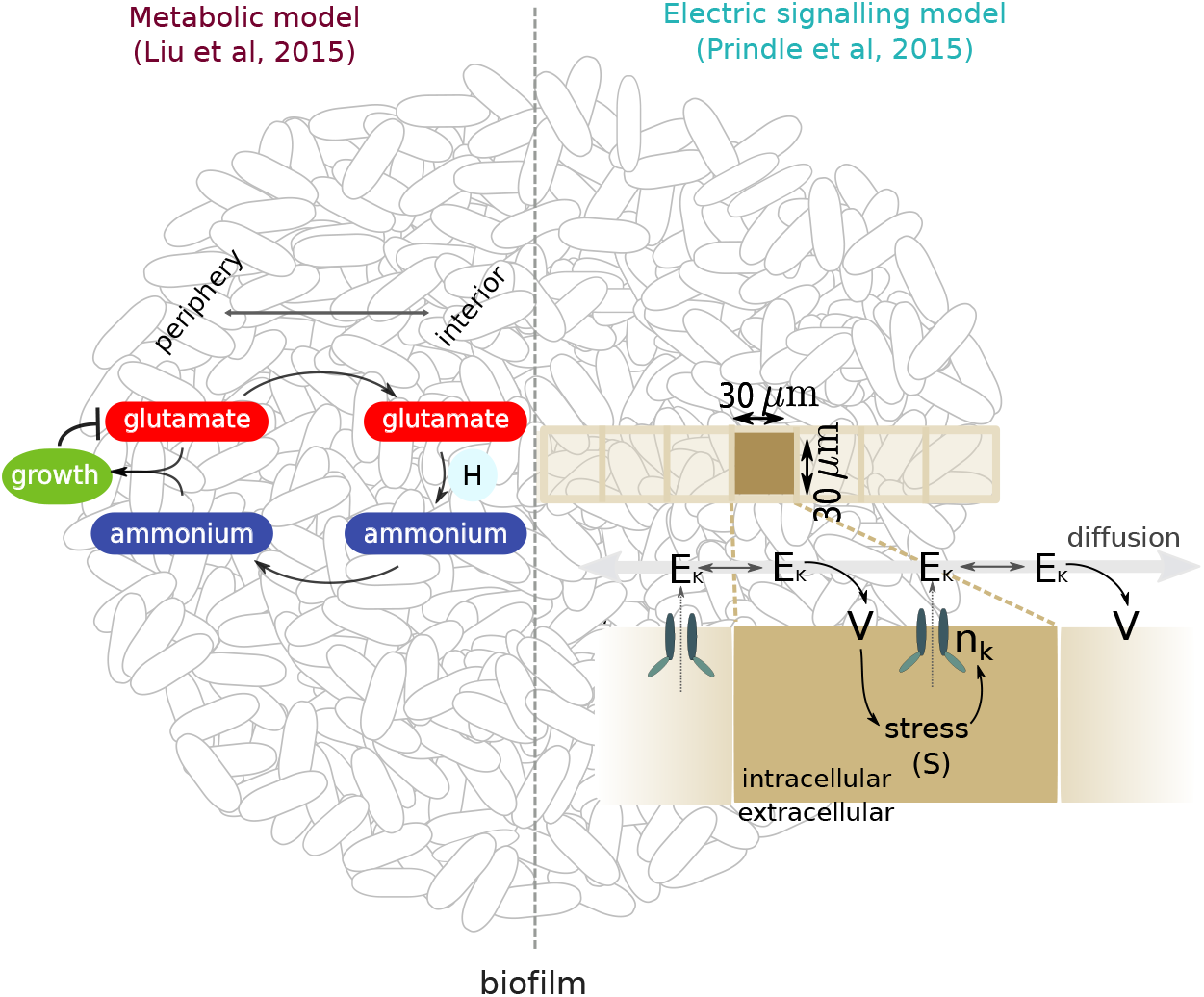
Metabolic (left) and electric (right) processes involved in the response of the biofilm to a gradient of nutrient-limitation-drive stress from the interior to the periphery. The left-hand side schematises the interactions considered in the metabolic model of Ref. [5], whereas the right-hand side schematises the electrical signalling propagation model proposed in Ref. [7]. In the diagram on the left, H represents the enzyme GDH. In the diagram on the right, E_K_ stands for excess extracellular potassium, V for the membrane potential, and n_k_ for the gating variable of the potassium channel.

Notably, potassium and glutamate are the two most abundant ions in living cells [8]. In addition to their their roles in osmoregulation, pH maintenance and nitrogen metabolism, they are essential for information propagation in animal nervous systems. In particular, potassium is one of the key ions involved in the modulation of the membrane potential in neurons, and glutamate is a central excitatory neurotransmitter. Besides the presence of the same ions in the two scenarios, there are also phenomenological links. In a normally functioning brain, electrical signalling typically propagates along neurons through action potentials mediated by sodium and potassium that can travel at speeds up to meters per second [9]. In addition, under pathological conditions, electrical activity can also propagate over the brain on a much slower timescale, on the order of millimetres per minutes (reviewed in Ref. [10]), similar to the biofilm potassium waves. This occurs during the phenomenon of cortical spreading depression, a wave of intense depolarisation followed by inhibition of electrical activity that entails a strong metabolic disturbance of neuronal function. Cortical spreading depression has been shown to occur in the brains of multiple species, from insects [11, 12] to mammals [13], where it has been regarded as an evolutionary conserved process [10] that has been linked to human pathologies like migraine, and to the propagation of brain damage during ischemic stroke. The phenomenon has been modelled as a reaction-diffusion process [14, 15] according to the following mechanism: an initial stimulus leads to an increase in extracellular potassium, causing the membrane potential to rise beyond a critical threshold that triggers a self-amplifying release of potassium, a major ionic redistribution of sodium, calcium and chloride, and release of neurotransmitters like glutamate. Diffusion of potassium and glutamate to neighbouring cells leads to subsequent depolarisation beyond a threshold, thus causing ionic release and self-propagation of the wave [10, 16, 17].

The widespread importance of potassium and glutamate, and the links between electrical signalling in bacteria and neuronal dynamics led us to investigate in more detail the roles of those two ions in the oscillations exhibited by *B. subtilis*. In this work, we unify in a discretised reaction-diffusion scheme the metabolic and electrical components of the system. Assuming a homogeneous population of cells, the model reveals the spontaneous emergence of two phenotypically-distinct populations of cells as a consequence of the spatiotemporal dynamics of the system, which causes oscillations to start beyond a critical size. We begin by developing a model that accommodates the interactions considered in previous works both for the metabolic and electrical components of the phenomenon. Then we show that the details of the ammonium metabolism are not required to explain the key aspects of the dynamics. The behaviour can be explained by a model with minimal interactions between glutamate metabolism and potassium signalling, thus clarifying the process in bacteria and providing further insight into the roles of these ubiquitous biological ions.

## 2 A unified spatially-extended model of biofilm oscillations

Conceptually, biofilm oscillations can be understood to arise as result of a delayed, spatially-extended negative feedback of metabolic stress [6]. A first molecular model was proposed in Ref. [5] (left half of Fig. 2), based on the fact that cells need to create glutamine from glutamate and ammonium for biomass production. Since ammonium is not present in the MSgg medium used in the experiments, it must be synthesised by the cells from glutamate. The model assumed that only cells in the interior of the biofilm produce ammonium (through the glutamate dehydrogenase enzyme GDH, H in Fig. 2), part of which diffuses away to the periphery of the biofilm, where it is used by the peripheral cells to grow. In turn, peripheral growth reduces glutamate availability in the centre, and subsequently ammonium availability in the periphery. As a result, peripheral cells stop growing, thus allowing the centre to recover glutamate and produce ammonium again, leading to a new oscillation cycle. The model considered two different cell populations –interior and peripheral–, and was subsequently implemented on a continuous spatial domain in Ref. [18], with still pre-defined interior and peripheral cell types.

This model can readily generate oscillations, which crucially rely on the assumption of a strongly nonlinear activation of GDH by glutamate in the interior population of cells (modelled with a Hill function with large Hill coefficient). However, it does not account for the electrical signalling component of the phenomenon, which we incorporate next.

### 2.1 Full model description

We consider a horizontal cross-section through the middle of a biofilm (as depicted in Fig. 1A) which allows us to simplify the system into a one-dimensional lattice, as shown in Fig. 3. The left-most lattice sites are ‘biofilm’ (shaded squares, ‘b’), followed by ‘non-biofilm’ sites (white squares, ‘n’) on the right. We begin by a more detailed model (Fig. 3) that includes all the aforementioned metabolic interactions [5] (left half of Fig. 2) in addition to the electrical signalling propagation [7] (right half of Fig. 2). The details of the equations can be found in the Supplementary, and we next describe the main points of the model (see also Table 1).

**Table 1:** Main assumptions and comparison between the full and simplified models.

| Model assumption | Full model<br>(section 2) | Simplified model<br>(section 3) |
| --- | --- | --- |
| Extracellular glutamate flow and diffusion | Yes | Yes |
| Ammonium flow and diffusion | Yes | No (no ammonium present) |
| Extracellular potassium flow and diffusion | Yes | Yes |
| Decay of flow and diffusion rates towards the biofilm interior as a consequence of cellular density | Yes | Yes |
| Glutamate uptake favoured by hyperpolarisation and disfavoured by depolarisation | Yes | Yes |
| Glutamate uptake enhanced in metabolically active cells | Through the effect of biomass-producing biomolecules | Directly linked to intracellular glutamate |
| Saturable glutamate uptake | Yes | Yes |
| GDH dynamics regulated by glutamate | Yes | No (no GDH present) |
| Ammonium produced from intracellular glutamate by GDH | Yes | No (no ammonium present) |
| Growth favored by high levels of intracellular glutamate | Through intermediate biomass-producing biomolecules. | Directly modulated by intracellular glutamate |
| Metabolic stress inhibited by intracellular glutamate | Yes | Yes |
| Potassium channel opening probability ($n_k$) increased by stress | Yes | Yes |
| Potassium released as a result of the opening probability ($n_k$) and membrane potential | Yes | Yes |
| Potassium uptake governed by homeostatic processes and the metabolic state of the cells (intracellular glutamate) | Yes | Yes |
| Membrane potential determined by intracellular and extracellular potassium concentrations | Yes | Yes |
| ThT as a reporter of membrane potential | Yes | Yes |

**Figure 3:**
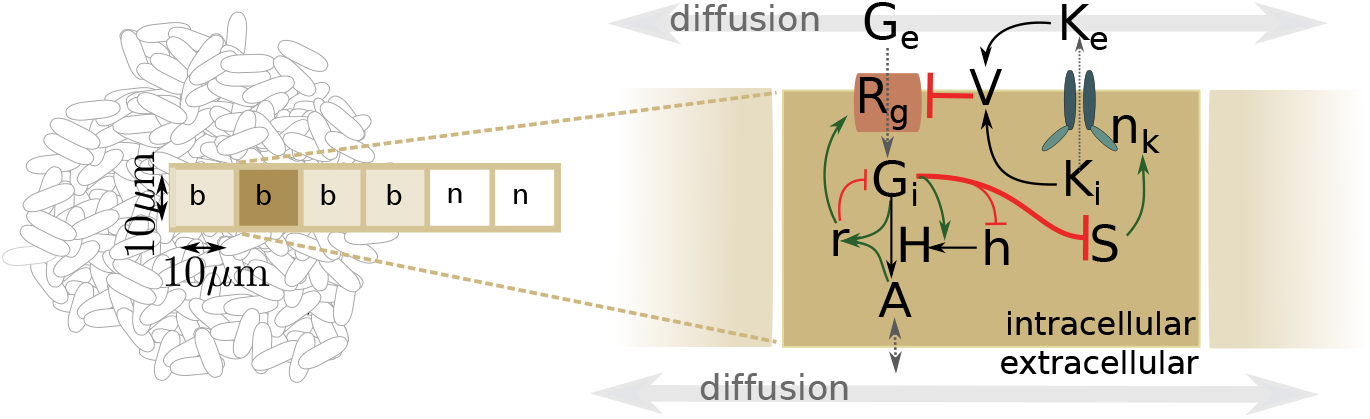
Scheme of the unified metabolic-electric model introduced in this work. In the lattice scheme on the left, the labels ‘b’ and ‘n’ inside the sites denote ‘biofilm’ and ‘non-biofilm’ lattice sites, respectively. In the network diagram on the right, G, K, h, H and A represent glutamate, potassium, inactive GDH, active GDH and ammonium, respectively. The quantity r represents biomass-producing biomolecules such as ribosomal proteins. R_g_ denotes the glutamate receptor, V the membrane potential, *n*_k_ the gating variable of the potassium channel, and S the stress. h stands for inactive GDH. The subindices ‘i’ and ‘e’ label intracellular and extracellular quantities, respectively. The two thick flat arrows highlight the links between the metabolic and electrical components of the system.

‘Non-biofilm’ sites have extracellular variables only: extracellular glutamate (*G*_*e*_), ammo-nium (*A*, assumed to have the same concentration inside and outside cells, following [5]), and extracellular potassium *K*_*e*_. These variables diffuse between neighboring lattice sites. In ad-dition, we also account for the fact that media is flowing constantly through the microfluidics chamber such that their concentration tends to become close to that in the media (Supplementary Eqs. (S1), (S2), (S17)).

‘Biofilm’ sites correspond to small biofilm regions, such that variable values would correspond to averages across multiple cells. As schematised in Fig. 3, in this sites there are also the extracellular variables (also subject to diffusion and media flow) in addition to the intracellular ones that take part in the various biochemical reactions. We model explicitly the dynamics of extracellular (*G*_*e*_) and intracellular (*G*_*i*_) glutamate in the biofilm (Supplementary Eqs. (S5)-(S6)), where we consider glutamate uptake into the cells and the conversion of glutamate into ammonium by the action of the enzyme GDH. In turn, we model GDH synthesis in its inactive form (*h*) and subsequent activation (*H*) (Supplementary Eqs. (S8)-(S9)). We assume that high concentrations of glutamate inhibit GDH synthesis. This follows from the original observation that GDH overexpression in the periphery kills biofilm oscillations [5], suggesting that GDH expression is restricted to the centre, where glutamate levels are low. Moreover, we assume that there is a threshold for GDH activation by glutamate, as in the original metabolic model. Glutamate and ammonium are assumed to be used for the production of biomass-producing biomolecules such as ribosomal proteins, denoted by *r* (Supplementary Eq. (S11)). We also account explicitly for the glutamate transporter concentration (*R*_*g*_), which we assume to saturate for large enough *G*_*e*_. In order to accommodate the assumption that cells with a higher metabolic activity consume more glutamate, we consider that *R*_*g*_ is subject to an inducible activation that depends on the presence of biomass-producing biomolecules (Supplementary Eq. (S7)).

In [7], the YugO potassium channel, which has a TrkA gating domain, was determined to be responsible for potassium release in this phenomenon. TrkA proteins are potassium channel regulatory proteins with nucleotide-binding domains that bind NAD/NADH [19, 20] and ADP/ATP [21], thus coupling potassium translocation to the metabolic state of the cell. Glutamate is a central amino acid at the cross-road between carbon and nitrogen metabolism. Our assumption is that low glutamate levels lead to imbalances in the aforementioned nucleotides that are sensed by TrkA and trigger potassium release [7]. Therefore, we assume that high levels of intracellular glutamate inhibit the production of stress-related biomolecules, resulting in the following equation for stress dynamics:

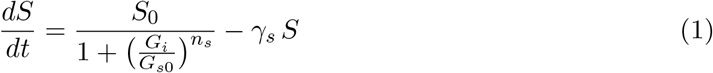

In turn, *S* enhances the opening probability of the potassium channel, given by *n*_*k*_. This results in a link from the metabolic component of the phenomenon to the electrical counterpart.

As the potassium channel opens, potassium is released to the extracellular environment. Conversely, potassium uptake is assumed to be governed by homeostatic processes that tend to keep its intracellular concentration at a fixed value, in addition to depend on the cellular metabolic state (glutamate level), to account for the energy demand of the process (Supplementary Eqs. (S13)-(S14)). Extracellular and intracellular potassium concentrations determine the membrane potential (*V*), whose dynamics is described by a Hodgkin-Huxley-like conductance-based model (Supplementary Eq. (S18)). Finally we include the ThT reporter 𝒯 downstream of the membrane potential, increasing when the cells become hyperpolarised due to potassium release (Supplementary Eq. (S20)).

In *B. subtilis*, glutamate is imported by pumps such as the symporter GltP [22], which uses the proton motive force as a source of energy, and this is in turn influenced by the membrane potential [23]. We therefore consider that glutamate transport into the cell is modulated by the membrane potential *V*, such that depolarisation reduces entry, and hyperpolarisation enhances it. This provides the link from the electrical to the metabolic component of the phenomenon, and is represented with the following switch-like import-modulation term:

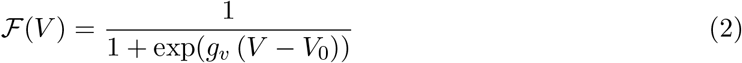

The biofilm is assumed to expand proportionally to the biomass-producing biomolecules. In order to simulate growth, we consider that ‘non-biofilm’ lattice sites neighbouring biofilm sites become occupied with cells (and thus become a lattice site of type ‘biofilm’) with probability *P*_*grow*_ *dt r*, with *dt* being the simulation time step. Importantly, each new biofilm lattice site inherits the intracellular variables of the ‘mother’ site. Due to this particularity, the continuum approximation is unsuitable, and we simulate the system as a coupled map lattice.

### 2.2 Simulations of the model

The diffusion coefficient in water for ions and small molecules such as potassium and glutamate is of the order of ∼ 10^6^*μ*m^2^/h. We thus fixed the diffusion coefficient of potassium and ammonium in the media to this value, and assumed the diffusion coefficient of glutamate to be half that value, due to its larger molecular weight. Regular glutamate concentration in the media is 30 mM and potassium concentration is 8 mM. Since there is no ammonium in the medium, *A*_*m*_ = 0. We fixed the resting membrane potential to *-*150 mV. Furthermore, oscillations experimentally start at a biofilm size of around 600 *μ*m with a period of around 2 hours [6], and potassium and ThT signals should be correlated [7]. With these constraints, we manually adjusted the rest of the parameters (Table S1) in order to reproduce the experimentally observed oscillatory dynamics.

Figure 4 shows kymographs for extracellular potassium and ThT (variables that can be experimentally monitored as explained above, see Fig. 1), as well as for extracellular glutamate, stress, active GDH and ammonium, which have not been experimentally quantified. Oscillations with a period of about two hours emerge in all variables once the biofilm becomes large enough. Moreover, in agreement with the experimental data, the top left plot shows that potassium peaks are likely to coincide with periods of no growth.

**Figure 4:**
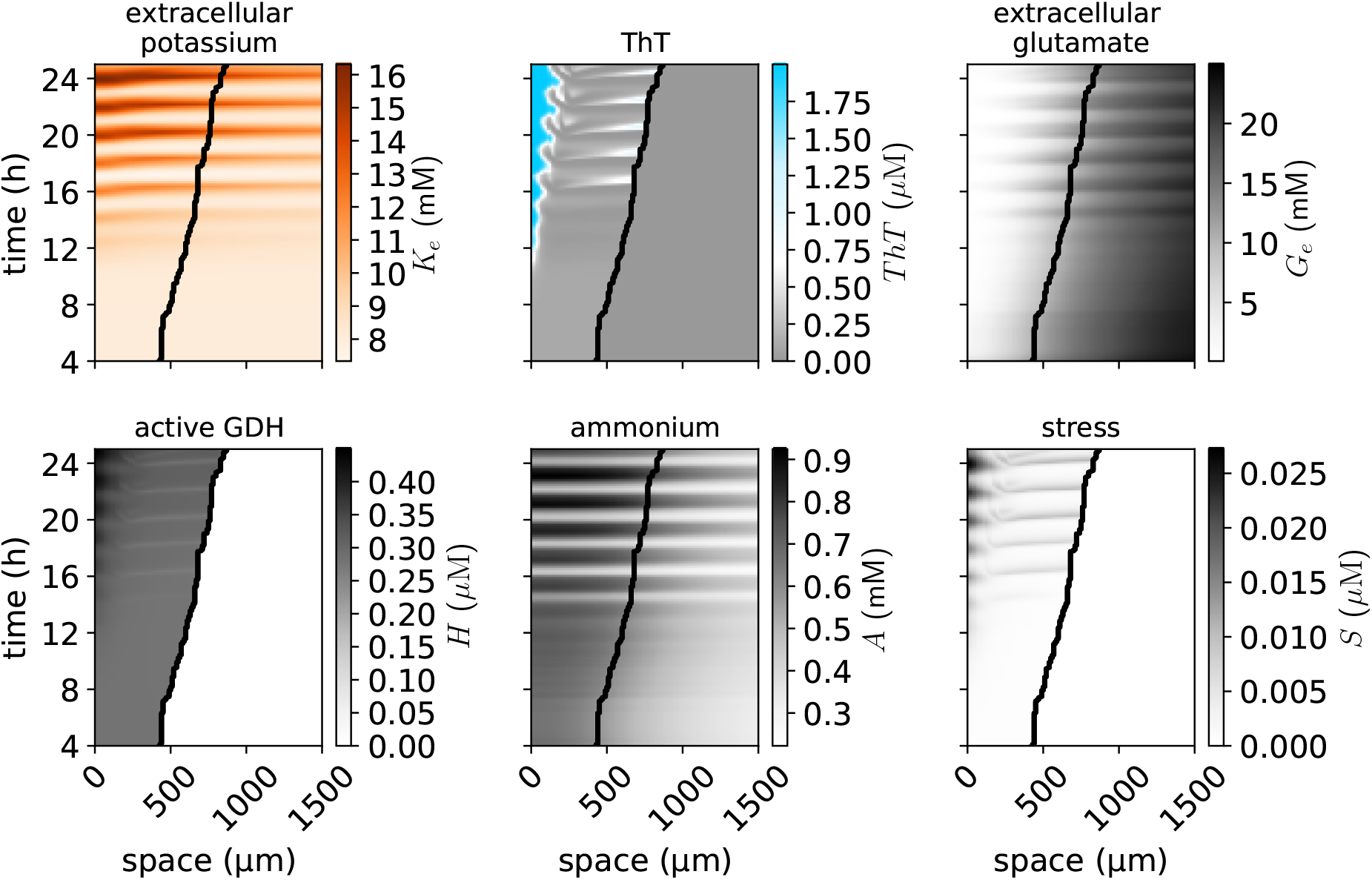
Oscillations in the full metabolic-electrophysiological model. Kymographs of the model simulation results, showing oscillations in various model variables.

In contrast to the original model of Ref. [5] (see also Ref. [18]), where two populations of cells (central and peripheral) were pre-defined regarding GDH production, in the current model an *a priori* separation of cell types is no longer required, but emerges spontaneously. This can be seen in the bottom left kymograph of Fig. 4, which shows that a central population with higher levels of GDH activity and stress emerges over time. Notably, this model does not require strong cooperativity in GDH activation for the oscillations to emerge (*n*_*H*_ = 2, in contrast to the value of 7 in Ref. [5] and 12 in Ref. [18]). This is likely to be the result of the electrical component of the model. In order to test the relevance of this component of the oscillations, we next simplify the metabolic details and consider only the interplay between glutamate metabolism and electrical signalling.

## 3 Oscillations emerge from the interplay between glutamate and electrical signalling

We now simplify the metabolic aspects of the full model introduced above, eliminating GDH, ammonium and biomass-producing biomolecules (Supplementary Eqs. (S8)-(S11)), and supposing instead that intracellular glutamate directly increases the glutamate transporter and directs growth (Fig. 5A). With this, the equations governing the metabolic part of the phenomenon become:

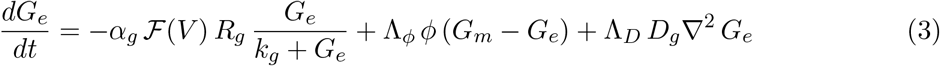

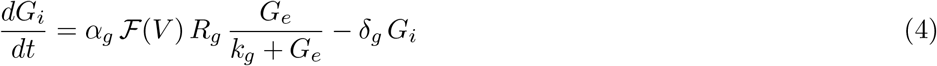

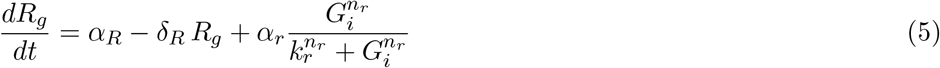

in addition to considering glutamate diffusion in the ‘non-biofilm’ sites (Supplementary Eq. S1).

**Figure 5:**
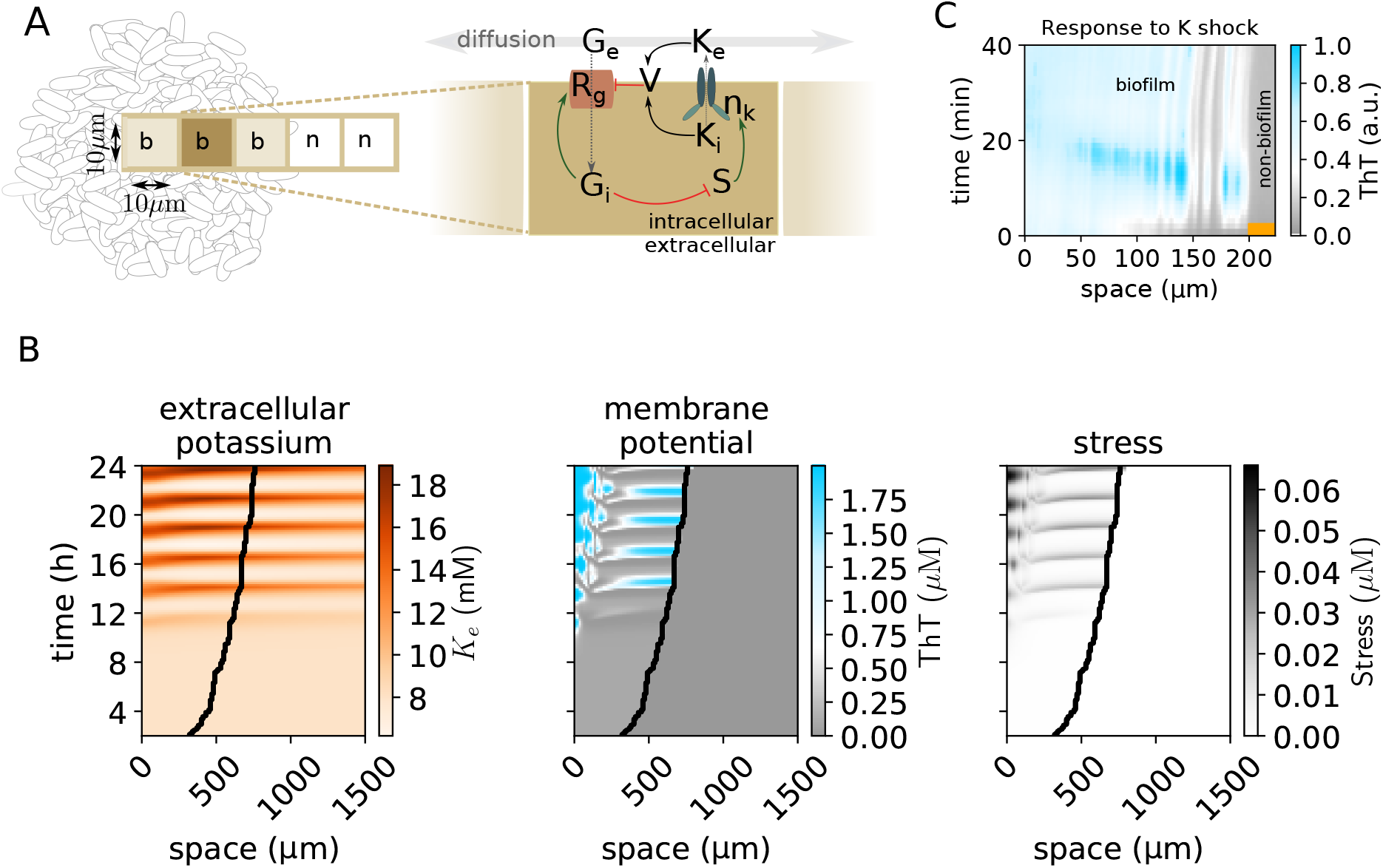
Oscillations in a simplified model, with glutamate and the electrical signalling compo-nents. A) Scheme of the model. B) Kymographs of various variables of the system. C) Electrical signalling wave (reported by ThT) in an experimental biofilm triggered by an increase of potassium in the media from 8 mM to 300 mM during 3 minutes (denoted by the orange mark). The wave propagates inwards, from the periphery (right) to the centre (left).

In this case, the probability with which the biofilm grows depends directly on the concentration of intracellular glutamate at the biofilm boundary according to *P*_*grow*_ *dt G*_*i*_. The dynamics of the electrical part of the model remain unchanged (Supplementary Eqs. (S12)-(S20)).

The kymographs in Fig. 5B show that oscillations beyond a critical size also emerge in this case. The results suggest the following mechanism for the oscillations: once the biofilm reaches a critical size, glutamate in the centre of the biofilm is too low and leads to metabolic stress. As a result, cells release potassium, that actively propagates towards the edge due to depolarisation and subsequent metabolic stress. Depolarisation leads to a drop in glutamate consumption levels in the peripheral region, due to the immediate effect on the transport efficiency, and a slightly delayed effect through the reduction in the glutamate transporter. As a result, glutamate can diffuse inwards to allow stress relief in the centre, and a new oscillation cycle can start.

According to this, oscillation onset requires stress in the centre to surpass a threshold that leads to potassium release. As we have recently described [6], oscillations can be triggered in experimental biofilms by transiently stopping the media flow of the microfluidics chamber. This is also the case in this model, where oscillations can be triggered by transiently setting the flow-rate parameter (*ϕ*) to zero. If the biofilm is sufficiently large, such a perturbation leads to a sudden reduction in the glutamate availability and stress increase, with the subsequent potassium release and oscillatory onset (Fig. S1).

Moreover, the model predicts that depolarising any region of the biofilm should be sufficient to trigger a self-propagating wave of stress and potassium release, with the associated changes in membrane potential. This is reproduced experimentally as shown in Fig. 5C: a short increase in the concentration of potassium in the media triggers a wave in experimental biofilms, which propagates inwards from the original (peripheral) depolarisation site.

## 4 Glutamate metabolism and stress release determine oscillation onset size and period

According to the aforementioned mechanism for the oscillations, if a biofilm increases its glutamate consumption, or if the glutamate concentration in the media is reduced, oscillations should start earlier because the centre becomes stressed at a smaller biofilm size. This expectation is fulfilled when simulations are performed with halved glutamate concentration in the media (*G*_*m*_), in good agreement with the experimental data (compare the first two conditions in the left panel of Fig. 6, see also Refs. [5, 6]). Similarly, bacteria that cannot synthesise glutamate due to a deletion in the *gltA* gene are expected to consume more glutamate from the media. We model this condition as an increase in both glutamate uptake rate (*α*_*g*_) and degradation rate (*δ*_*g*_), as a result of higher glutamate demand. Also in this case oscillations start earlier both in the model and the experiments (third condition in the left panel of Fig. 6).

**Figure 6:**
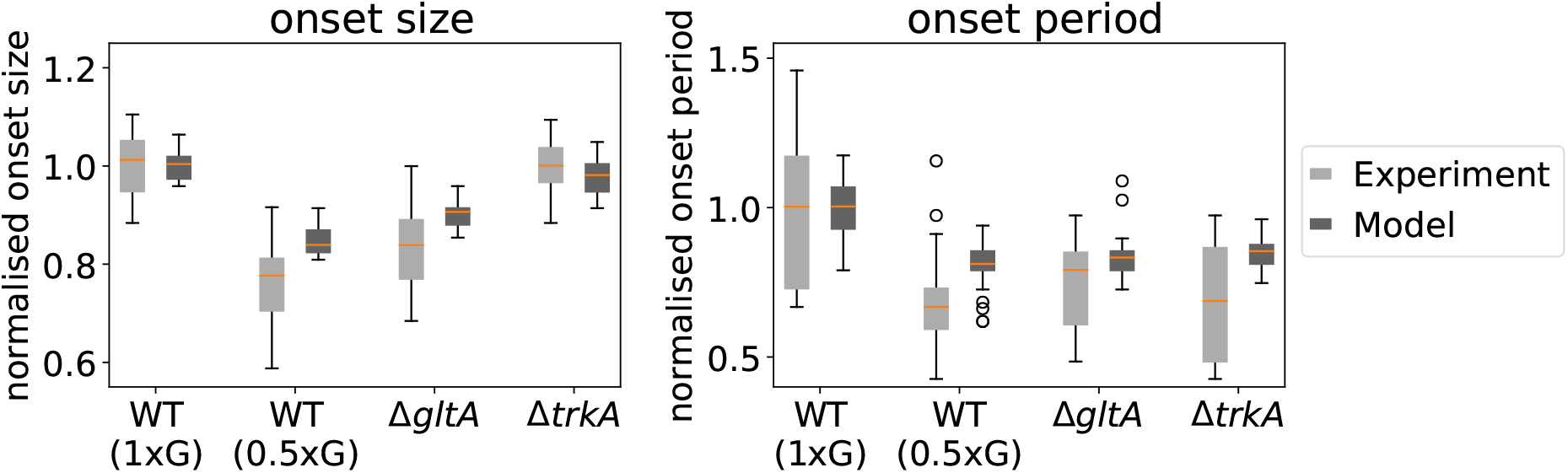
Biofilm size and oscillation period at onset. Light grey: Experimental Data. Dark grey: Model. The data is normalised with respect to the average of each WT condition (model and experiments separately). Boxplots extend between the first and third quartiles of the normalised data, with a line (orange) at the median. The extreme of the whiskers denote the range of the data within 1.5x the interquartile range, with outliers plotted individually. Experimental data contains a minimum of 12 data points per condition. See Methods for details on the simulations.

In these two cases of reduced onset size, the onset period is also smaller than in the reference situation (first three conditions in the right panel of Fig. 6). As we have described in previous work [6], larger biofilms tend to have higher periods, the explanation being that the larger a biofilm is, the more time it takes for the stress signal to propagate from centre to periphery, and thus the longer the oscillation period.

However, according to the oscillation mechanism that this model suggests, the period of the oscillations must also depend on the metabolism dynamics and the stress relieving capabilites of the cells due to the potassium effects. The Δ*trkA* mutant strain was characterised in Ref. [7] to be deficient in electrochemical signalling, due to the deletion of the gating domain of the YugO channel. The mutation can be interpreted to render a channel with reduced conductance and increased basal leak [7]. This can be modelled with increased *a*_0_ (to simulate a leaky channel), and reduced *g*_*K*_ (to simulate reduced conductance once the channel is open). The model simulations predict that oscillations start at the same size as with the basal parameters, but with a reduced period, in good agreement with the experimental data (last condition in Fig. 6). The unaffected onset size is consistent with the fact that glutamate metabolism is largely unperturbed in this mutant, whereas the altered potassium channel leads to less effective stress relief during wave propagation (Figs. S2), and thus facilitates a subsequent cycle. Therefore, the relationship between biofilm size and oscillation period is not universal, but depends upon the metabolic and electrical signalling capabilities of the cells and thus on the genetic background of the strain.

## 5 Discussion

The oscillations in growth rate and membrane potential exhibited by *B. subtilis* biofilms is an instance of self-organising, emergent behaviour similar to the complex collective dynamics that characterizes neuronal tissue. Here we have proposed a unified conceptual framework for these bacterial oscillations that combines glutamate metabolism with potassium wave propagation. Our discrete reaction-diffusion scheme assumes an initially homogeneous population of cells, and exhibits a spontaneous emergence of oscillations beyond a critical biofilm size. Our work shows that oscillations are triggered by a decrease in glutamate levels in the biofilm centre, that lead to metabolic stress and potassium release. Potassium diffusion to the neighbors interferes with glutamate metabolism, thus causing a self-propagating wave of membrane potential changes that allows glutamate recovery in the centre and thus explains the oscillation cycle.

With some notable recent exceptions in multicellular eukaryotes [24], electrophysiology has only been considered relevant so far for excitable cells such as those in cardiac and neural tissue. The resemblance, both at the molecular and functional level, between excitable wave propagation in these tissues and the communication of metabolic stress among bacterial cells calls for an evolutionary perspective on the phenomenon of electrical signalling. Animal nervous systems have evolved intricate electrochemical circuits that allow sensing and responding to both internal and external stimuli, with high integrative and computational capabilities clearly evidenced by the human brain. How this complexity arose is still a matter of active investigation [25, 26, 27, 28] but it is tempting to speculate that the bacterial oscillations reported here may represent an ancient instance of electrically-mediated information transmission.

Primitive prokaryotic potassium channels are regarded as the ancestors of the animal cation channels whose diversification has been linked to the evolution of nervous systems [29, 30]. Given that high intracellular concentrations of potassium are essential for bacterial pH maintenance [31, 8], it is natural to expect that potassium channels were used in early bacteria for osmoregulation [27]. The biofilm oscillations studied here suggest that potassium also acts as a signalling intermediate in modern bacteria, communicating information among cells of both the same and also different bacterial species [32].

The link between metabolism and electrical signalling described here is also reminiscent of the relationship between cellular energy and neuronal function [33]. Neuronal dynamics is highly dependent upon cellular metabolism and energy state: the high energy requirements of the ion pumps that maintain proper electrochemical gradients are well known [34]. In addition, some animal voltage-gated ion channels directly respond to energy-related metabolites like NAD(H) and ADP/ATP through direct ligand binding [35], as in the bacterial YugO channel.

While the mechanisms that allow information processing in bacterial biofilms share similar chemical and electrical processes with animal brains, the latter compute at much faster speeds and thus in much more complex ways. This can be partly ascribed to the reliance of animal brains on faster voltage-gated ion channels, such as those dependent on sodium, which evolved only when rapid responses were increasingly beneficial (due for instance to the appearance of predators) [27]. Nevertheless, slow propagation of electrical activity is still present in modern animal brains. During the phenomenon of cortical spreading depression, for instance, a slow wave of electrical activity propagates over brain tissue, with massive ionic redistribution involving, critically, potassium and glutamate accompanied by a strong metabolic disturbance. This phenomenon has been regarded as intrinsic to neurons and is evolutionarily conserved across animal species [10]. In the light of these facts, it is tempting to see the potassium-mediated transmission of stress among bacterial cells as a precursor of more recent electrically-mediated information propagation tasks in animal brains. The study of membrane potential dynamics and electrical signalling processes in other bacterial species will be important to further illuminate this issue.

## Methods

### 5.1 Experimental data

#### Strains and Plasmids

All experiments were performed using *Bacillus subtilis* strain NCIB 3610. For the Δ*trkA* mutant, we deleted the C-terminal portion of *yugO* (amino acids 117328), leaving only the N-terminal ion channel portion of YugO (amino acids 1116). For the Δ*gltA* mutant, the *gltA* gene was replaced by kanamycin resistant gene.

#### Growth conditions

Biofilms were grown using the standard MSgg biofilm-forming medium [36]. In this media glutamate is the only nitrogen source for the bacteria. We explored biofilm dynamics at glutamate concentrations of 1× (30 mM) and 0.5× (15 mM). For microfluidics, We used the CellASIC ONIX Microfluidic Platform and the Y04D microfluidic plate (EMD Millipore). Details can be found in our previous work [5, 37].

#### Time-Lapse Microscopy

Biofilms were monitored using time-lapse microscopy, using Olympus IX81 and IX83 inverted microscopes. To image entire biofilms, 10× lens objectives were used. Images were taken every 10 min. We tracked membrane potential dynamics using the fluorescent dyes Thioflavin T (10 *μ*M) and APG-4 (2*μ*M), from TEFLabs [7].

### 5.2 Modeling

#### Simulation methods

Simulations were performed using custom-code in C, and analysis and plotting was done in Python. We used a first-order approximation of the discrete Laplacian operator [38] and the time evolution of the variables was obtained using a 4th order Runge-Kutta method. Boundary conditions were reflective: the outside neighbours of the lattice sites at the boundary of the system are the boundary sites themselves (such that the discrete normal derivatives of the variables at the boundary are zero). The simulation time step was set to 5 × 10^-6^ h, and each lattice site corresponds to 10 *μ*m. The system was allowed to relax towards the steady state at the beginning of the simulation by preventing expansion during 1 hour of simulated time. The initial conditions for the intracellular variables are *S* = 0 *μM*, *n*_*k*_ = 0, *V* = *-*156 mM, *K*_*i*_ = 300 mM, *ThT* = 0 *μM*, *Rg* = 1 *μM*, *H*_*i*_ = 0.5, *H* = 0.25, *R* = 1, and *G*_*i*_ is log-normally distributed around 3 mM. In the biofilm cells, *K*_*e*_ is log-normally distributed around 8 mM, and *A* = 1 mM. In the non-biofilm cells, *A* = 0. Initial *G*_*e*_ = 30 mM (or 15 mM in reduced glutamate concentrations).

#### Numerical perturbations

In order to study the effect of halved glutamate in the media, and the *trkA* and *gltA* deletions, we assumed the following parameter values with respect to the WT: Δ*gltA* mutant: *δ*_*g*_ = 1.5×, *α*_*gt*_ = 1.3×; Δ*trkA* mutant: *a*_0_ = 2×, *g*_*K*_ = 0.5×. We performed 30 simulations per condition starting from a random initial radius between 150 and 250 *μm*. Potassium peaks were defined based on a threshold of 9.5 mM (pulse begins when crossing the threshold). Onset period is defined to be the time between the beginnings of the first two peaks, and onset size that of the first beginning.

## SUPPLEMENTARY MATERIAL

### 1 Model equations and description

As explained in the main text, we consider a one-dimensional array of simulation lattice sites where biochemical species react and diffuse. Here we describe in detail the model equations.

#### 1.1 Metabolic component

In the ‘non-biofilm’ sites, we consider media flow and diffusion of glutamate (*G*) and ammonium (*A*) according to the following equations:

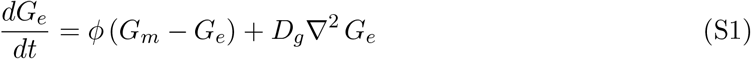

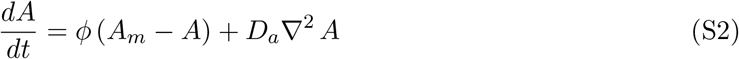

In the microfluidics chamber, media is flowing constantly. Thus, we simulate the effect of the media flow with the first term of each equation, such that with some rate *ϕ*, the concentration of the chemical species tends to equate that in the medium (*X*_*m*_). The second term models diffusion.

In the biofilm, we assume that the diffusion coefficient and flow rate of glutamate decay exponentially with the distance to the biofilm edge *d*_*e*_, due to the extracellular matrix and high cell density, according to the following functions:

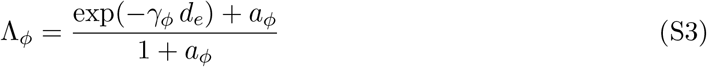

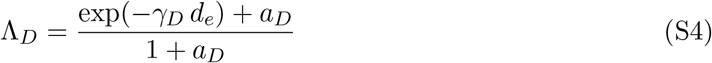

These functional forms are chosen such that at the edge Λ_*x*_ = 1, and flow and diffusion match those in the media. The coefficients decay towards the biofilm interior, tending asymptotically to *a/*(1 + *a*) in the centre, which ensures some remaining flow and diffusion. We follow the original assumption [5] that ammonium diffusion over the biofilm is very fast, and do not apply these reduction terms to this variable.

The dynamics of extracellular (*G*_*e*_) and intracellular (*G*_*i*_) glutamate in the biofilm are modelled with the following dynamical equations:

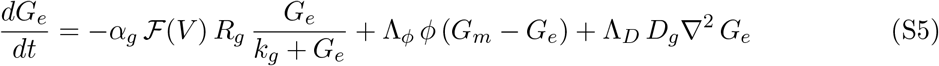

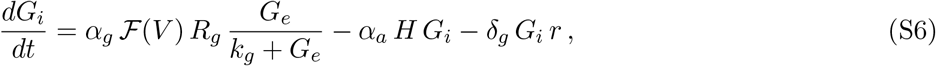

where, as mentioned, *G*_*e*_ is affected by media flow and diffusion. The first term in the right-hand side of the two equations represents glutamate transport into the cells. As explained in the main text, we consider that glutamate transport into the cell is modulated by the membrane potential *V*, such that depolarisation reduces entry, and hyperpolarisation enhances it, according to the functional form given in Eq. (2) of the main text.

In addition, we assume that glutamate is imported into the cells through the glutamate transporter *R*_*g*_, which saturates for large enough *G*_*e*_, with half-maximum concentration *k*_*g*_. We describe explicitly the dynamics of *R*_*g*_ by:

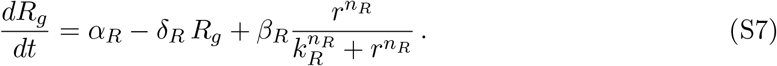

We thus assume that *R*_*g*_ is produced at a basal rate *α*_*R*_ and degraded at a rate *δ*_*R*_. The last term accounts for the higher glutamate uptake by metabolically active cells, such that the presence of biomass-producing biomolecules, such as ribosomal proteins, denoted by *r*, enhances *R*_*g*_ synthesis via a Hill function with exponent *n*_*R*_.

Equation (S6) also assumes that intracellular glutamate concentration decays due to ammonium production via the GDH enzyme (represented by *H* in the *α*_*a*_-term at the right-hand side of the equation) and through various metabolic tasks including in particular biomass production (*δ*_*g*_-term).

The production of ammonium is regulated by the activity of the enzyme GDH. We describe the dynamics of the inactive and active forms of this enzyme, *h* and *H* respectively, by the equations:

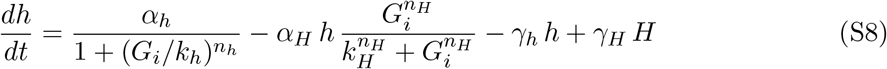

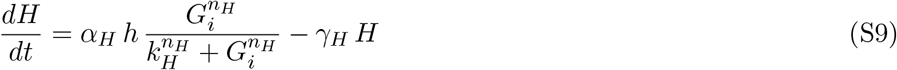

such that we account for synthesis and degradation of inactive GDH and its conversion into the active form (*α*_*H*_-term in the two equations). As explained in the main text, we assume that high concentrations of glutamate inhibit GDH synthesis, whereas activation is positively regulated by glutamate via a Hill function with exponent *n*_*H*_. We also consider deactivation at a constant rate *γ*_*H*_.

Ammonium dynamics is affected by production from glutamate by active GDH, consumption for various metabolic processes such as biomass production, and diffusion:

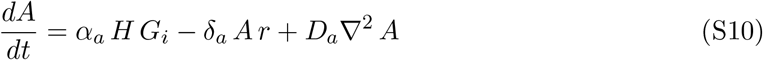

Finally, biomass production is considered to increase with ammonium and intracellular glutamate, and to be subject to linear decay:

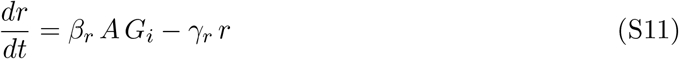

#### 1.2 Electrical signalling

Next we incorporate an adapted version of the electrical model introduced in [7]. As explained in the main text, we consider an inhibitory effect of intracellular glutamate on a stress variable *S*, whose production rate is modelled with an inhibitory Hill function:

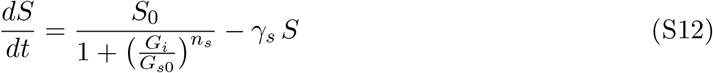

We explicitly consider both extracellular (*K*_*e*_) and intracellular (*K*_*i*_) potassium, whose dynamics are given by:

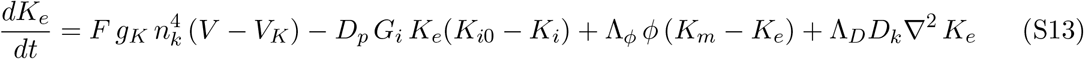

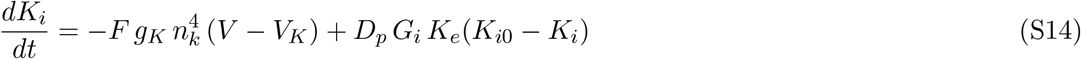

Potassium uptake is assumed to be governed by homeostatic processes that tend to keep its intracellular concentration at a fixed value *K*_*i*0_, described by the second term in the right-hand side of Eqs. (S13)-(S14). Uptake is also made to depend on the metabolic state (glutamate level), to account for the energy demand of the process. In addition, extracellular potassium diffuses and is subject to the media flow in the chamber.

Potassium flow through its ion channel (first term in the right-hand side of the *K*_*e*_ and *K*_*i*_ equations) is governed by the corresponding Nernst potential:

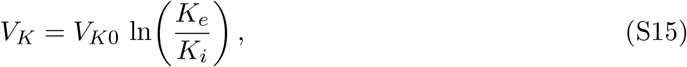

and depends on the opening probability of the potassium channel, *n*_*k*_:

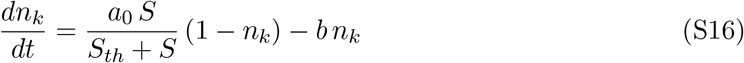

In the media lattice sites, as in the case of glutamate and ammonium [Eqs. (S1) and (S2)], the extracellular potassium dynamics is affected by diffusion and by the media flow:

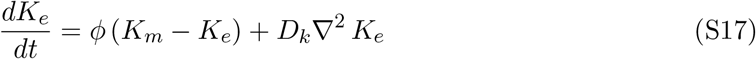

The membrane potential dynamics is described by a Hodgkin-Huxley-like conductance-based model containing potassium flux through the ion channel and a leak current [7]:

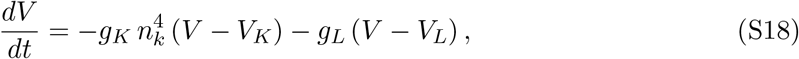

where the leak potential *V*_*L*_ is assumed to depend on the extracellular potassium [7] in a threshold-linear manner, such that when *K*_*e*_ is larger than its basal level in the medium, *K*_*m*_, the leak potential *V*_*L*_ grows linearly (and the cell depolarizes), while when *K*_*e*_ *< K*_*m*_ the leak potential stays at its basal level:

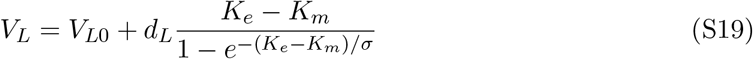

Finally, we include the ThT reporter 𝒯 downstream of the membrane potential, increasing when the cells become hyperpolarised due to potassium release:

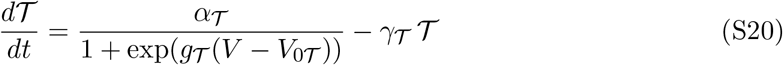

#### 1.3 Simplified model

In the simplified model, we keep the same dynamics for the electrical part (Eqs. (S12)-(S20)). The metabolic part is simplified as follows: Eqs. (S8)-(S11) are removed, the equations for extracellular glutamate dynamics, in both ‘biofilm’ and ‘non-biofilm’ sites (Eqs. S1, S5), are maintained, and Eqs. (S6) and (S7) become:

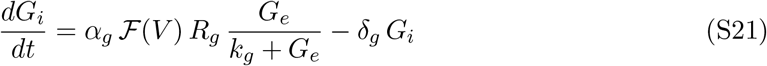

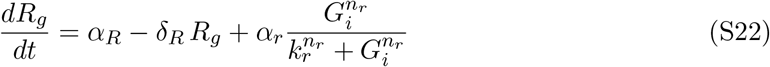

### 2 Supplementary tables and figures

**Table S1:**
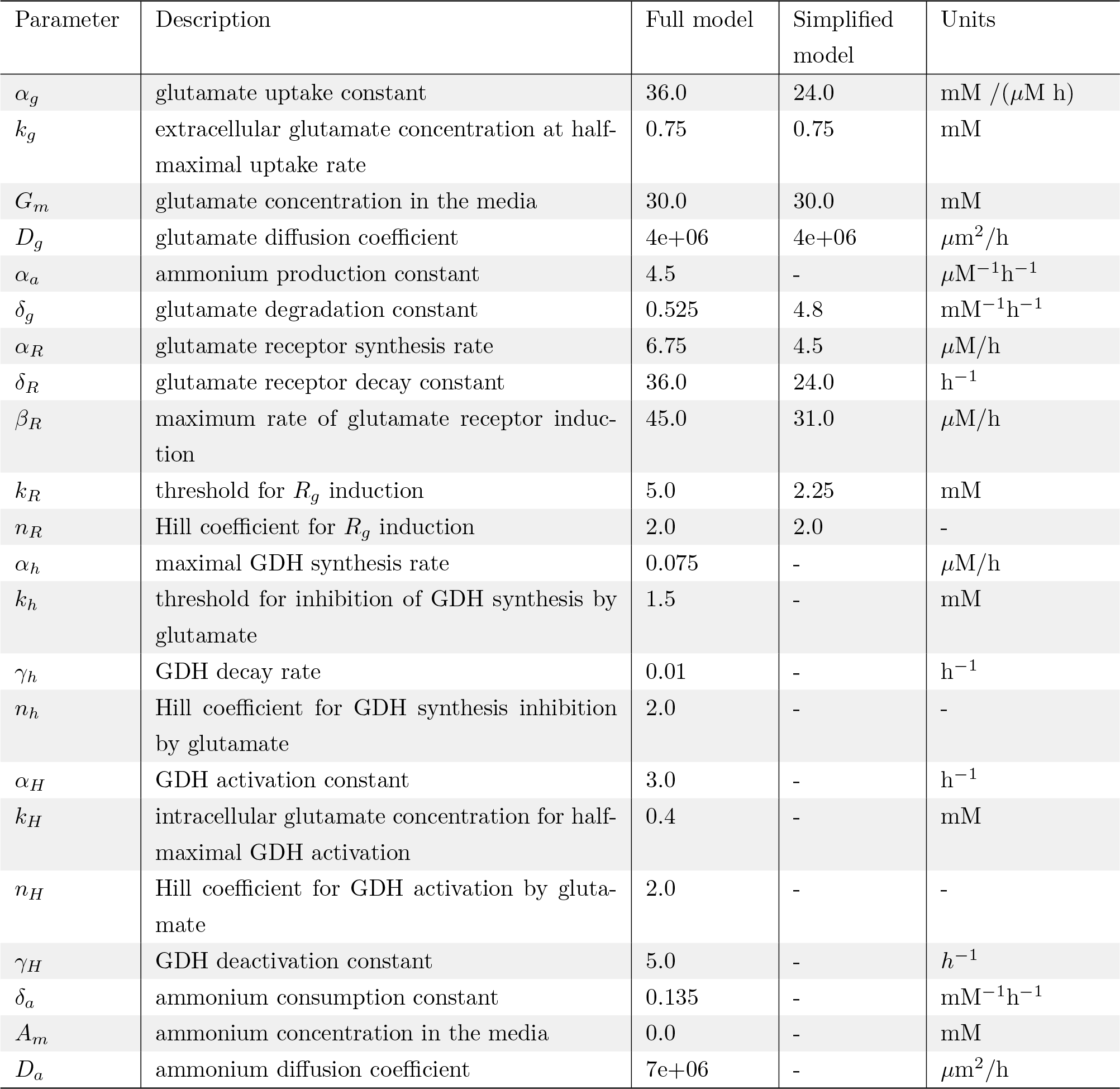
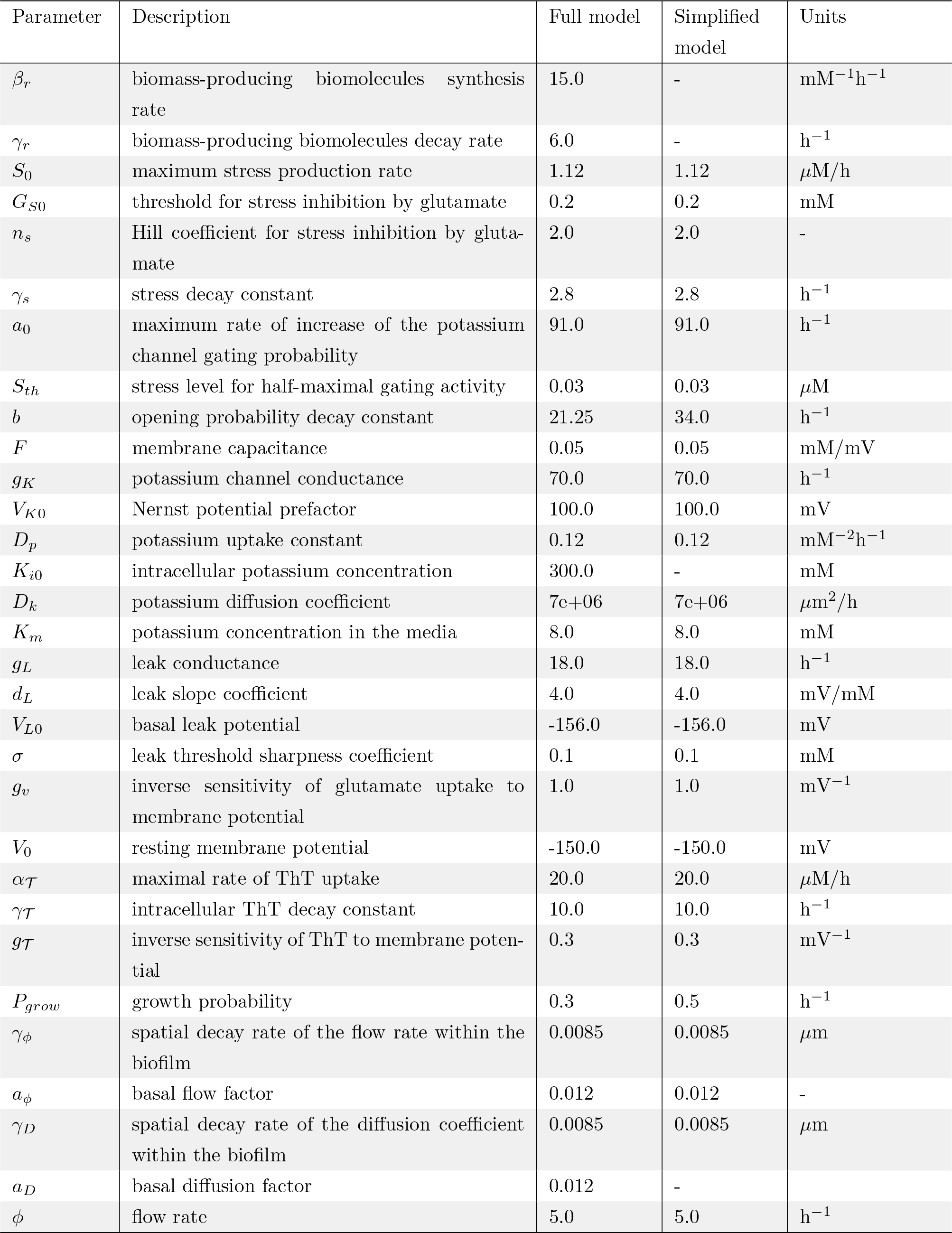
Parameter description and basal values for the two models.

**Figure S1:**
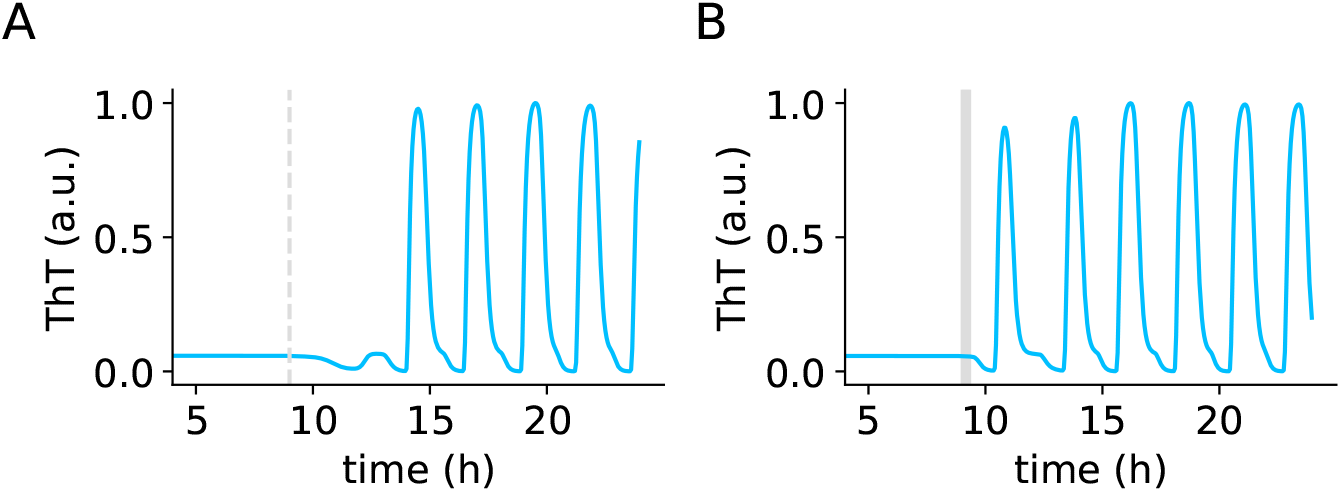
Stop-flow triggers oscillations in the simplified model. Normalised ThT time traces at the periphery (50 *μ*m from the biofilm edge). A) Reference simulation. Vertical dashed line indicates the time of stop-flow in B. (This simulation is the same as in Fig. 5B). B) The biofilm was perturbed with a stop-flow-like perturbation (*ϕ* = 0 during 20 min at *t* = 9 hours). The growth noise realisation is the same in both simulations, such that the only difference is the perturbation.

**Figure S2:**
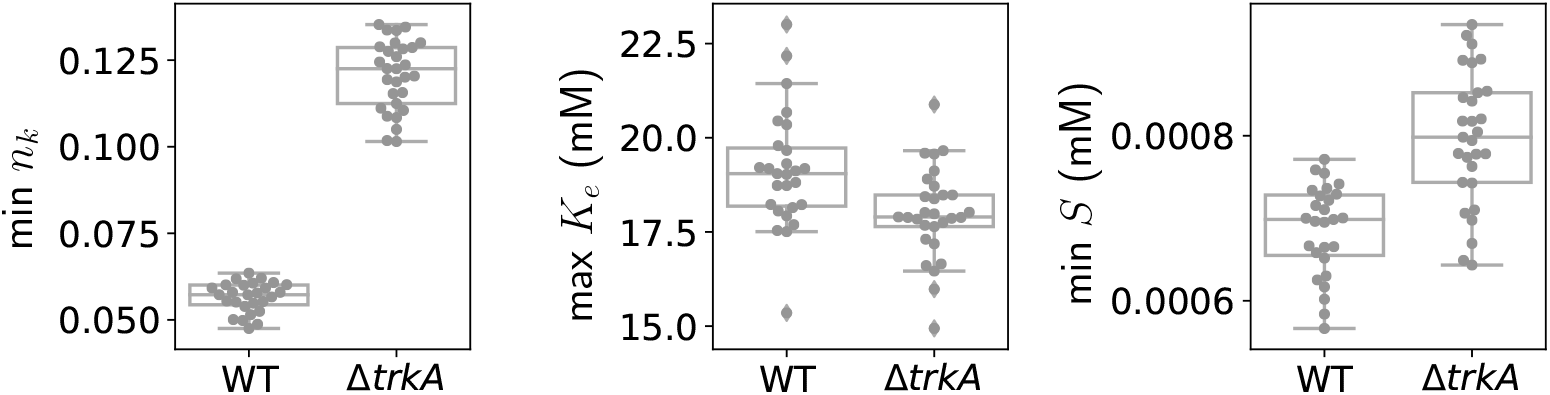
The Δ*trkA* mutation in the model leads to impaired stress relief. For each simulation, either the maximum or the minimum of the variable during the oscillations was computed for the peripheral region (outermost 100 *μ*m). Each dot represents a simulation, from the same data as in Fig. 6 from the main text.

